# Magnetotactic bacteria accumulate a large pool of iron distinct from their magnetite crystals

**DOI:** 10.1101/2020.03.10.986679

**Authors:** Matthieu Amor, Alejandro Ceballos, Juan Wan, Christian P. Simon, Allegra T. Aron, Christopher J. Chang, Frances Hellman, Arash Komeili

## Abstract

Magnetotactic bacteria (MTB) are ubiquitous aquatic microorganisms that form intracellular nanoparticles of magnetite (Fe_3_O_4_) or greigite (Fe_3_S_4_) in a genetically controlled manner. Magnetite and greigite synthesis requires MTB to transport a large amount of iron from the environment which is subsequently concentrated in organelles called magnetosomes for crystal precipitation and maturation. X-ray absorption analysis of MTB suggests that the intracellular iron is mainly contained within the crystals, thus preventing potential toxic effects of free iron. In contrast, recent mass spectrometry studies suggest that MTB may contain a large amount of iron that is not precipitated in crystals. Here, we attempt to resolve these descrepancies by performing chemical and magnetic assays to quantify the different iron pools in the magnetite-forming strain *Magnetospirillum magneticum* AMB-1 cultivated at varying iron concentrations. AMB-1 mutants showing defects in crystal precipitation were also characterized following the same approach. All results show that magnetite represents at most 30 % of the total intracellular iron under our experimental conditions. We further examined the iron speciation and subcellular localization in AMB-1 using the fluorescent indicator FIP-1 that is designed for detection of labile Fe(II). Staining with this probe suggests that unmineralized reduced iron is found in the cytoplasm and associated with magnetosomes. Our results demonstrate that, under our experimental conditions, AMB-1 is able to accumulate a large pool of iron distinct from magnetite. Finally, we discuss the biochemical and geochemical implications of these results.

**Importance:** Magnetotactic bacteria (MTB) are a group of microorganisms producing iron-based intracellular magnetic crystals. They represent a model system for studying iron homeostasis and biomineralization in bacteria. MTB contain an important mass of iron, about 10 to 100 higher than other bacterial model such as *Escherichia coli*, suggesting efficient iron uptake and storage systems. Accordingly, MTB have been proposed to significantly impact the iron biogeochemical cycle in sequestering a large amount of soluble iron into crystals. Recently, several studies proposed that MTB could also accumulate iron in a reservoir distinct from their crystals. Here, we present a chemical and magnetic methodology for quantifying the fraction of the total cellular iron contained in the magnetic crystals of the magnetotactic strain *Magnetospirillum magneticum* AMB-1. Comparison of the mass of iron contained in the different cellular pools showed that most of the bacterial iron is not contained in AMB-1 crystals. We then adapted protocols for the fluorescent detection of Fe(II) in bacteria, and showed that iron could be detected outside of crystals using fluorescence assays. This work suggests a more complex picture for iron homeostasis in MTB than previously thought. Because iron speciation controls its solubility, our results also provide important insights into the geochemical impact of MTB. A large pool of unmineralized iron in MTB could be more easily released in the environment than magnetite, thus limiting iron sequestration into MTB crystals.

## Introduction

Many living organisms transform inorganic molecules into crystalline structures in a process called biomineralization. Magnetotactic bacteria (MTB) represent an elegant example of such organisms. They incorporate dissolved iron from their environment and precipitate it as magnetite [Fe(II)Fe(III)_2_O_4_] or greigite [Fe(II)Fe(III)_2_S_4_] nanoparticles in organelles called magnetosomes (Uebe and Schüler, 2016). MTB are ubiquitous gram-negative microorganisms in aquatic environments. They inhabit the oxic/anoxic transition zones in the water column or sediments where they thrive (Kopp and Kirschvink, 2008). In MTB, magnetosomes are aligned as chains inside the cell, and provide the bacteria with a permanent magnetic dipole presumably for navigation purposes (Uebe and Schüler, 2016).

Tremendous work has been carried out to determine the biological and chemical reactions leading to magnetite synthesis in MTB (McCausland and Komeili, 2020). In the two best-studied, magnetite-forming strains *Magnetospirillum magneticum* AMB-1 and *Magnetospirillum gryphiswaldense* MSR-1, magnetosome formation is a genetically-controlled process where: (*i*) magnetosome vesicles are formed from invagination of the inner cell membrane, (*ii*) empty magnetosome vesicles are aligned as a chain inside the cell, (*iii*) iron is transported and concentrated into magnetosome for initiation of biomineralization, and (*iv*) crystal size and shape are precisely controlled in a species-specific manner. A set of ∼30 genes, located in a distinct portion of the genome called the magnetosome island (MAI), is required and sufficient for the step-wise formation of magnetosomes (Murat *et al*, 2010). Recently, iron isotope studies have provided an integrative model for iron uptake and precipitation as magnetite in the magnetotactic strain AMB-1 (Amor *et al*, 2016, 2018). This model assumes that dissolved Fe(II) or Fe(III) species are incorporated into AMB-1 and stored in the cytoplasm and/or periplasm as Fe(III). This Fe(III) pool is then partially reduced into Fe(II) for trafficking to magnetosomes, and oxidized for precipitation as magnetite (Amor *et al*, 2018). Direct mass spectrometry measurements of iron content and iron isotope composition in AMB-1 cells devoid of magnetosomes suggests that a large pool of iron, which could represent at least 50% of the total cellular iron, accumulate in reservoir(s) distinct from magnetite (Amor *et al*, 2016, 2018). The above-mentioned high-resolution mass spectrometry measurements of iron showed discrepancy with previous X-ray absorption analyses performed on both AMB-1 and MSR-1 strains, in which magnetite was the sole iron species detected at the end of the biomineralization process (Baumgartner *et al*, 2013; Fdez-Gubieda *et al*, 2013).

In the present work, we address the discrepancy between the mass spectrometry and X-ray absorption experiments by determining the distribution of iron in AMB-1 cells. We grew AMB-1 with different iron concentrations, and measured the mass of iron taken up by the bacteria using chemical assays. Cells were then recovered and the mass of iron contained in magnetite was quantified from magnetic characterizations. From these experimental results, we found that magnetite represents ∼25 to ∼30% of the bulk cellular iron. Additional cultures of AMB-1 were grown to determine the total mass of iron contained in magnetite at the scale of a bacterial population. Simultaneous cell counting allowed us to estimate the mean mass of magnetite per cell. Comparison of these results with published single-cell quantification of bulk iron in AMB-1 supported mass balance estimations. Finally, we used a fluorescent reporter of iron to show that at least part of the non-crystalline iron is present as Fe(II) species within bacterial cells. To further investigate the link between magnetite formation and iron incorporation, mutant AMB-1 strains (Δ*mamP* and Δ*mamT* strains lacking the protein MamP and MamT, respectively) showing biomineralization defects were also analyzed following the same approaches (Fig. 1). Bacterial cultures of all strains were carried out in triplicates.

**FIG 1.**
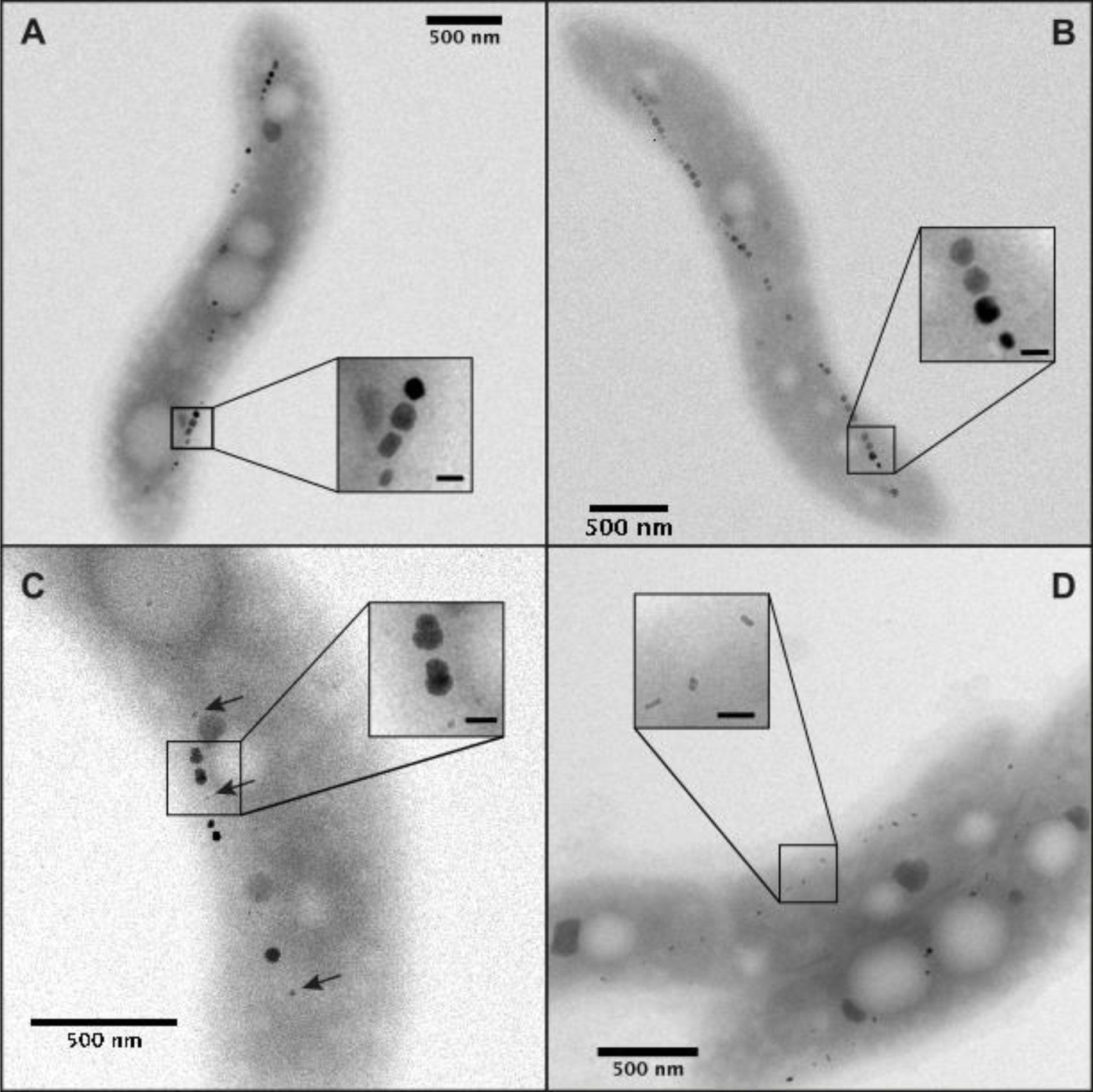
Transmission electron observations of wild-type AMB-1 cells cultivated for three days with initial iron concentrations in the growth medium of (A) 30 or (B) 150 μM, and (C) Δ*mamP* and (D) Δ*mamTΔR9* (referred to as Δ*mamT*) AMB-1 strains cultivated with Fe(III)-citrate at 150 μM. Arrows in (C) indicate small nanoparticles produced by the Δ*mamP* bacteria. Scale bars (insets) = 50 nm.

All data support the presence a large pool of iron, at least partially reduced, distinct from magnetite in AMB-1 under our experimental conditions. These results raise important biochemical (*i.e.* iron homeostasis in MTB) and geochemical (*i.e.* impact of MTB on the iron biogeochemical cycle) questions that we address in the discussion of this article.

## Results

### Iron depletion and speciation in AMB-1 cultures

We first cultivated wild-type AMB-1 (Fig. 1) for three days at two iron concentrations (30 and 150 μM). Sterile media containing no bacteria were also prepared and used as a control condition. The iron concentration and oxidation state were monitored in AMB-1 cultures and sterile media using the ferrozine assay (see materials and methods). Iron concentration and oxidation state in the filtered sterile media remained constant over the three days of incubation, showing that all iron in the growth media can be analyzed by the ferrozine assay (Table 1; Fig. 2). Therefore, the decrease in iron concentration in the growth media is not due to precipitation of small iron phases excluded during filtration and can be attributed to iron uptake by AMB-1. Initial and final Fe(II), total iron concentration and pH in wild-type AMB-1 growth media, as well as final optical densities (OD_400nm_), are given in Table 2. Most of the bacterial iron uptake, normalized to biomass, occurred between one and two days of culture (Fig. 2): iron depletion was 0.10 ± 0.04 and 0.95 ± 0.21 mg per unit of optical density after 26 hours of culture for an initial iron concentration of 30 and 150 μM, respectively; and 0.37 ± 0.03 and 1.05 ± 0.15 mg per unit of optical density after 46 hours of culture for an initial iron concentration of 30 and 150 μM, respectively (Fig. 3A). For an initial concentration of 30 μM, no further iron depletion occurred after 46 h of culture. In contrast, the mass of iron depleted from the growth medium decreased to 0.69 ± 0.18 mg per unit of optical density in cultures provided with 150 μM of iron. Iron speciation was also modified over bacterial growth. 40 and 17% of the initial Fe(III) added to the growth media containing 30 and 150 μM of iron, respectively, became immediately reduced after inoculation (Fig. 4). Remaining Fe(III) was then progressively reduced until complete reduction, which happened after 46 and 69 hours of culture for an initial iron concentration of 30 and 150 μM, respectively. Finally, the growth medium pH showed a similar increase between the two iron conditions, with final values of ∼7.5 (Table 2).

**FIG 2.**
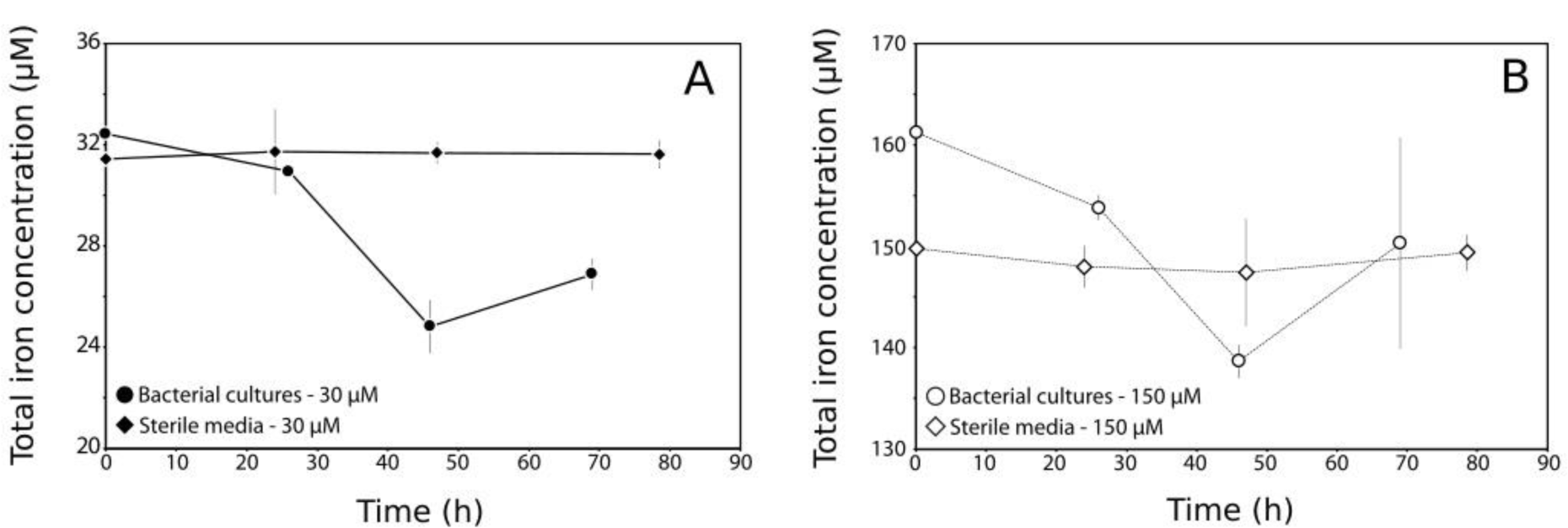
Total iron concentration in (circles) AMB-1 and (diamonds) sterile media provided with iron at (A) 30 or (B) 150 μM. Each point corresponds to the mean value of 3 replicates ± 1SD. Note the different y-axes.

**FIG 3.**
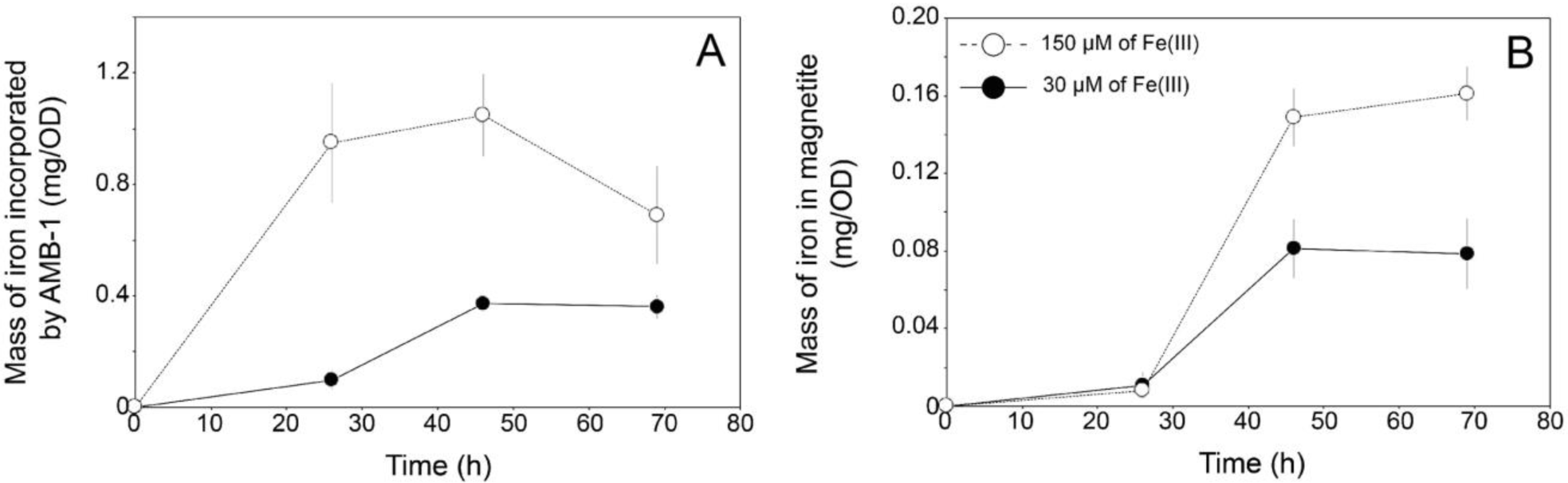
Mass of iron (A) taken up by AMB-1 and (B) contained in magnetite during bacterial growth. All values are normalized to optical densities (OD), which is proportional to the cell biomass. Thus, different cell densities cannot explain discrepancies in iron uptake. Each point corresponds to the mean value of three replicates ± 1SD. Note the different y-axes. Black circles and open symbols refer to cultures with an initial iron concentration of 30 and 150 µM, respectively.

**FIG 4.**
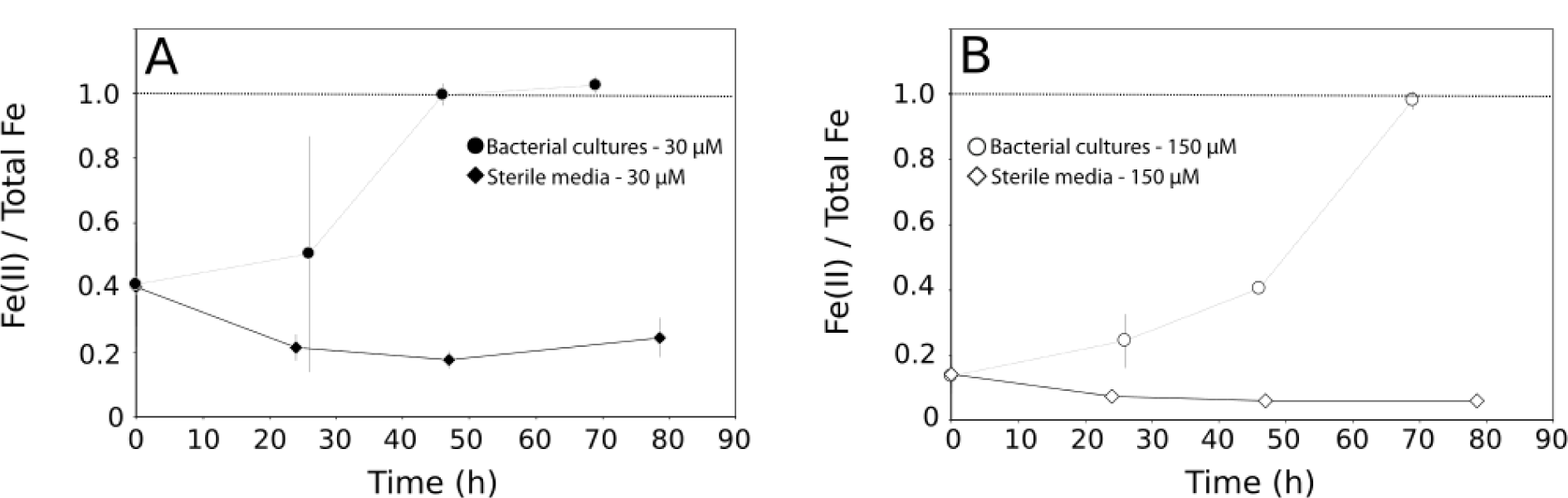
Iron speciation in (circles) AMB-1 and (diamonds) sterile media provided with iron at (A) 30 or (B) 150 μM. Each point corresponds to the mean value of 3 replicates ± 1SD.

**Table 1.** Fe(II) and total iron concentration in initial and final sterile growth media.

| Time of culture (h) | Volume of culture (mL) | Initial [Fe(II)] (μM) | Initial [Fe] (μM) | Final [Fe(II)] (μM) | Final [Fe] (μM) |
| --- | --- | --- | --- | --- | --- |
| **[Fe] = 30 μM** |  |  |  |  |  |
| 24 #1 | 190 | 21.41 | 33.07 | 8.53 | 33.59 |
| 24 #2 | 190 | 18.97 | 31.50 | 6.61 | 30.28 |
| 24 #3 | 190 | 18.27 | 32.02 | 5.39 | 31.33 |
| 47 #1 | 190 | 14.44 | 30.46 | 6.70 | 32.10 |
| 47 #2 | 190 | 12.18 | 32.20 | 5.29 | 31.22 |
| 47 #3 | 190 | 11.14 | 30.46 | 4.94 | 31.75 |
| 78.5 #1 | 190 | 8.70 | 31.15 | 9.88 | 31.22 |
| 78.5 #2 | 190 | 5.22 | 30.98 | 6.88 | 32.28 |
| 78.5 #3 | 190 | 4.35 | 31.15 | 6.53 | 31.39 |
| **[Fe] = 150 μM** |  |  |  |  |  |
| 24 #1 | 190 | 33.07 | 151.58 | 11.83 | 146.53 |
| 24 #2 | 190 | 29.59 | 151.06 | 10.44 | 149.49 |
| 24 #3 | 190 | 30.63 | 150.71 | - | - |
| 47 #1 | 190 | 24.02 | 152.45 | 10.93 | 153.27 |
| 47 #2 | 190 | 22.28 | 139.57 | 8.64 | 146.39 |
| 47 #3 | 190 | 19.14 | 151.41 | 7.05 | 142.68 |
| 78.5 #1 | 190 | 12.53 | 153.49 | 10.41 | 147.80 |
| 78.5 #2 | 190 | 11.66 | 149.49 | 9.70 | 149.03 |
| 78.5 #3 | 190 | 8.70 | 148.45 | 7.58 | 151.33 |

**Table 2.** Fe(II) and total iron concentration in the growth medium before and after AMB-1 cultures, and final optical density at 400 nm (OD_400nm_) and pH values of bacterial cultures (starting pH = 6.9).

| Time of culture (h) | Volume of culture (mL) | Initial [Fe(II)] (μM) | Initial [Fe] (μM) | Final OD<sub>400nm</sub> (AU) | Final pH | Final [Fe(II)] (μM) | Final [Fe] (μM) |
| --- | --- | --- | --- | --- | --- | --- | --- |
| **[Fe] = 30 μM** |  |  |  |  |  |  |  |
| 26 #1 | 185 | 16.87 | 31.36 | 0.067 | 6.95 | 7.29 | 30.94 |
| 26 #2 | 185 | 17.24 | 31.72 | 0.060 | 6.91 | 11.03 | 30.94 |
| 26 #3 | 185 | 17.05 | 31.72 | 0.083 | 7.01 | 28.45 | 30.94 |
| 46 #1 | 185 | 14.12 | 32.46 | 0.209 | 7.26 | 24.99 | 24.30 |
| 46 #2 | 185 | 13.94 | 32.82 | 0.202 | 7.34 | 24.99 | 26.03 |
| 46 #3 | 185 | 14.85 | 31.72 | 0.214 | 7.26 | 24.12 | 24.12 |
| 69 #1 | 185 | 8.80 | 33.19 | 0.212 | 7.42 | 28.12 | 27.42 |
| 69 #2 | 185 | 8.44 | 33.19 | 0.203 | 7.44 | 28.47 | 27.07 |
| 69 #3 | 185 | 7.52 | 33.74 | 0.204 | 7.42 | 26.20 | 26.20 |
| **[Fe] = 150 μM** |  |  |  |  |  |  |  |
| 26 #1 | 185 | 29.16 | 162.65 | 0.069 | 7.03 | 51.57 | 154.89 |
| 26 #2 | 185 | 26.96 | 157.15 | 0.067 | 6.94 | 35.21 | 152.40 |
| 26 #3 | 185 | 26.41 | 159.35 | 0.056 | 6.94 | 25.96 | 154.18 |
| 46 #1 | 185 | 26.77 | 161.00 | 0.215 | 7.18 | 57.80 | 140.24 |
| 46 #2 | 185 | 23.29 | 158.25 | 0.215 | 7.28 | 54.15 | 138.85 |
| 46 #3 | 185 | 20.90 | 161.73 | 0.211 | 7.30 | 56.58 | 136.94 |
| 69 #1 | 185 | 15.95 | 160.27 | 0.197 | 7.48 | 143.39 | 146.88 |
| 69 #2 | 175 | 15.95 | 172.00 | 0.202 | 7.50 | 154.57 | 162.08 |
| 69 #3 | 185 | 14.12 | 158.80 | 0.202 | 7.50 | 143.74 | 141.99 |

Iron uptake by the two mutant Δ*mamP* and Δ*mamT* strains was then measured following the same experimental procedure. Because mutant strains produce less magnetite than wild-type AMB-1, as shown by the electron microscope observations (Fig. 1), we cultivated them with iron at 150 μM to measure iron uptake more accurately. The highest iron uptake by wild-type AMB-1 was observed at ∼45 h of culture (see results). Accordingly, Δ*mamP* and Δ*mamT* strains were cultivated for ∼45 hours. Wild-type AMB-1 cultures were used as controls. The pH and optical density (OD) values were almost identical in all cultures of the three wild-type control, Δ*mamP* and Δ*mamT* AMB-1 strains (Table 3). Δ*mamP* cells showed limited iron incorporation normalized to the optical density, with a ∼4-fold decrease compared to the wild-type (Table 3). Iron uptake by Δ*mamT* cells showed inconsistent values, with bulk iron concentration showing a slight increase during bacterial growth in two of the three replicates, and a decrease in the third replicate. We considered these data to be inconclusive. Finally, the Fe(II) / total Fe ratios were slightly lower in Δ*mamP* (0.11 ± 0.02) and Δ*mamT* (0.09 ± 0.02) compared to wild-type AMB-1 (0.16 ± 0.03).

**Table 3.** Fe(II) and total iron concentration in the growth medium before and after mutant AMB-1 cultures, and final optical density at 400 nm (OD_400nm_) and pH values of bacterial cultures (starting pH = 6.9). Additional wild-type cultures were used as a control condition.

| Time of culture (h) | Volume of culture (mL) | Initial [Fe(II)] (μM) | Initial [Fe] (μM) | Final OD<sub>400nm</sub> (AU) | Final pH | Final [Fe(II)] (μM) | Final [Fe] (μM) |
| --- | --- | --- | --- | --- | --- | --- | --- |
| **Wild-type** |  |  |  |  |  |  |  |
| 43 #1 | 178 | 25.08 | 150.30 | 0.250 | 7.30 | 26.23 | 145.78 |
| 43 #2 | 173 | 22.85 | 155.69 | 0.252 | 7.24 | 25.23 | 150.59 |
| 43 #3 | 167 | 23.22 | 164.05 | 0.255 | 7.40 | 19.42 | 151.99 |
| ***ΔmamP*** |  |  |  |  |  |  |  |
| 43 #1 | 171 | 21.55 | 159.59 | 0.261 | 7.32 | 21.43 | 158.20 |
| 43 #2 | 174 | 19.32 | 161.27 | 0.252 | 7.35 | 18.23 | 161.00 |
| 43 #3 | 170 | 18.21 | 162.75 | 0.261 | 7.33 | 14.42 | 159.20 |
| ***ΔmamT*** |  |  |  |  |  |  |  |
| 43 #1 | 175 | 16.72 | 157.92 | 0.248 | 7.34 | 15.02 | 158.20 |
| 43 #2 | 177 | 15.42 | 153.23 | 0.280 | 7.35 | 15.62 | 148.29 |
| 43 #3 | 164 | 14.31 | 169.44 | 0.252 | 7.39 | 10.21 | 172.22 |

### Magnetic properties of AMB-1 cultures

After chemical analyses, bacteria were recovered and transferred into sample holders for magnetic analyses. Hysteresis loops were measured on whole bacterial populations (Fig. S1). Sample preparation was performed under anoxic conditions to prevent magnetite oxidation into maghemite [γ-Fe_2_O_3_]. Three magnetic parameters were extracted from hysteresis loops: the remanent magnetization (M_rs_), the saturation magnetization (M_s_) and the coercivity (H_c_). M_s_ depends only on the mass of magnetic material for a given phase, and is 92 emu per gram of magnetite (Zaitsev *et al*, 1999). Iron phases in MTB have been extensively described. In AMB-1 and MSR-1, as well as in the closely-related strain *Magnetospirillum magnetotacticum* MS-1, three iron species were evidenced both from bulk measurements and observations at the atomic scale: magnetite, ferrihydrite and Fe^2+^ (Frankel et al, 1983; Faivre et al, 2007; Baumgartner et al, 2013; Fdez-Gubieda et al, 2013; Uebe et al, 2019). Ferrihydrite and Fe^2+^ are paramagnetic at room-temperature, and do not to contribute to the M_s_ signal (Towe and Bardley, 1967; Aharoni, 2000; Wang et al, 2016). The M_s_ values thus provide accurate estimates of the mass of iron in magnetite, which is a ferrimagnetic material. M_rs_ corresponds to the remanent sample magnetization measured under an external magnetic field of zero, after exposing the sample to a saturating external field. M_rs_/M_s_ ratio depends on particle size and organization, and is typically ranging between 0.43 and 0.50 in AMB-1 (Dunlop, 2002; Li *et al*, 2012). Finally, H_c_ is the magnetic field strength required to reduce the magnetization of the sample to zero after fully magnetizing it. Thus, H_c_ represents the capability of a magnetic material to resist demagnetization. It depends on the particle size, its shape, its magnetization, and its magnetocrystalline anisotropy.

Magnetic parameters calculated from hysteresis loops (Fig. S1) are given in Table 4 and Fig. 5. After 26 hours of growth, remanent (M_rs_) and saturation (M_s_) magnetizations were similar between the two experimental conditions (Figs. 5A and 5B). Similar to iron uptake patterns, most magnetite was precipitated between 26 and 46 hours of culturing in both conditions. No variation in M_rs_ and M_s_ was observed for longer time of culture. Final M_rs_ was 9.57 ± 2.41 x 10^-4^ emu and 2.09 ± 0.57 x 10^-3^ emu for AMB-1 cultivated with iron at 30 and 150 μM, respectively. Final M_s_ values were 2.06 ± 0.44 x 10^-3^ and 4.13 ± 0.38 x 10^-3^ emu for AMB-1 cultivated with iron at 30 and 150 μM, respectively. Knowing the magnetic moment of magnetite per unit of mass (92 emu/g), the M_s_ values were converted to a mass of iron contained in magnetite. Results are given in Fig 3B. The maximum mass of iron in magnetite measured was 0.082 ± 0.015 and 0.15 ± 0.015 mg per unit of optical density in AMB-1 cultivated with 30 and 150 μM of iron, respectively. From the remanent and saturation magnetization values, we calculated the M_rs_ / M_s_ ratios. Almost identical values between the two experimental conditions were observed: ∼0.38, ∼0.50 and ∼0.50 after 26, 46 and 69 hours of cultures, respectively (Fig. 5C). Coercivity showed a slightly different pattern, as it progressively increased with time (Fig. 5D). After 26 hours of growth, the coercivity of AMB-1 cultures was ∼50 Oe for both initial iron concentrations. At longer growth times, AMB-1 cultivated with 150 μM showed higher coercivities (180 ± 5 Oe and 224 ± 31 Oe for 46 and 69 hours of growth, respectively) than AMB-1 cultivated with 30 μM of iron (132 ± 8 Oe and 146 ± 26 Oe for 46 and 69 hours of growth, respectively).

**FIG 5.**
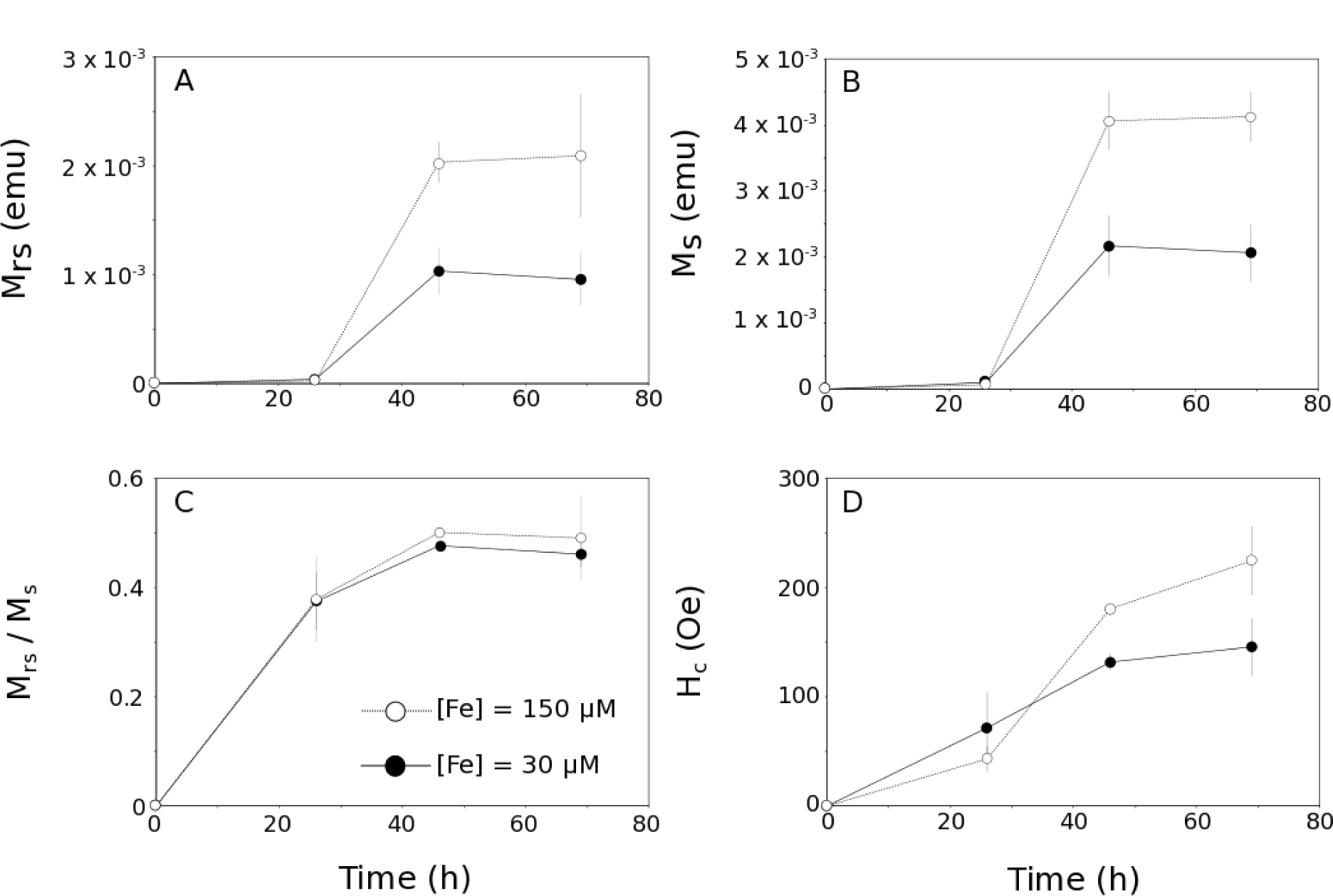
(A) Remanent magnetization (M_rs_), (B) saturation magnetization (M_s_), (C) coercivity (H_c_) and (D) M_rs_ / M_s_ ratios for the studied AMB-1 cultures. Each point corresponds to the mean value of three replicates ± 1SD. Black circles and open symbols refer to cultures with an initial iron concentration of 30 and 150 µM, respectively.

**Table 4.** Remanent magnetization (M_rs_), saturation magnetization (M_s_), coercivity (H_c_) and M_rs_/M_s_ ratios of wild-type AMB-1.

| Time of culture (h) | $M_{rs}$<br>(emu) | $M_s$<br>(emu) | $H_c$<br>(Oe) | $M_{rs}/M_s$ |
| --- | --- | --- | --- | --- |
| **[Fe] = 30 μM** |  |  |  |  |
| 26 #1 | $1.74 \times 10^{-5}$ | $4.06 \times 10^{-5}$ | 75.50 | 0.43 |
| 26 #2 | $2.55 \times 10^{-5}$ | $7.88 \times 10^{-5}$ | 37.02 | 0.32 |
| 26 #3 | $7.03 \times 10^{-5}$ | $1.89 \times 10^{-4}$ | 102.17 | 0.37 |
| 46 #1 | $1.08 \times 10^{-3}$ | $2.23 \times 10^{-3}$ | 138.02 | 0.48 |
| 46 #2 | $8.00 \times 10^{-4}$ | $1.68 \times 10^{-3}$ | 123.28 | 0.48 |
| 46 #3 | $1.22 \times 10^{-3}$ | $2.60 \times 10^{-3}$ | 134.23 | 0.47 |
| 69 #1 | $6.80 \times 10^{-4}$ | $1.57 \times 10^{-3}$ | 115.97 | 0.43 |
| 69 #2 | $1.07 \times 10^{-3}$ | $2.24 \times 10^{-3}$ | 156.12 | 0.48 |
| 69 #3 | $1.12 \times 10^{-3}$ | $2.39 \times 10^{-3}$ | 164.70 | 0.47 |
| **[Fe] = 150 μM** |  |  |  |  |
| 26 #1 | $5.19 \times 10^{-5}$ | $1.12 \times 10^{-4}$ | 56.81 | 0.46 |
| 26 #2 | $1.99 \times 10^{-5}$ | $6.42 \times 10^{-5}$ | 34.87 | 0.31 |
| 26 #3 | $1.15 \times 10^{-5}$ | $3.18 \times 10^{-5}$ | 37.50 | 0.36 |
| 46 #1 | $2.23 \times 10^{-3}$ | $4.52 \times 10^{-3}$ | 186.19 | 0.49 |
| 46 #2 | $2.01 \times 10^{-3}$ | $4.03 \times 10^{-3}$ | 175.58 | 0.50 |
| 46 #3 | $1.86 \times 10^{-3}$ | $3.64 \times 10^{-3}$ | 179.27 | 0.51 |
| 69 #1 | - | - | - | - |
| 69 #2 | $1.69 \times 10^{-3}$ | $3.89 \times 10^{-3}$ | 202.27 | 0.43 |
| 69 #3 | $2.49 \times 10^{-3}$ | $4.57 \times 10^{-3}$ | 246.70 | 0.55 |

We measured the mass of ferrimagnetic material in mutant AMB-1 strains following the same approach. Jones and co-workers showed that nanoparticles produced in the *ΔmamP* and *ΔmamT* cells also correspond to magnetite (Jones *et al*, 2015). Therefore, the same saturation magnetization per unit of mass (92 emu/g) was used to calculate the mass of iron contained in magnetite. Mutant AMB-1 strains showed altered magnetic properties (Table 5). The remanent and saturation magnetizations in the Δ*mamP* strain were ∼1 order of magnitude lower than wild-type. Associated M_rs_ / M_s_ ratios showed lower values in Δ*mamP* AMB-1 (0.34 ± 0.09) compared to the wild-type control (0.46 ± 0.05). In *ΔmamT* AMB-1, the saturation magnetization showed even lower values (∼2 orders of magnitude lower than the wild-type strain), while the remanent magnetization was ∼0. Accordingly, M_rs_ / M_s_ ratios corresponding to magnetite produced by Δ*mamT* cells were also ∼0. Finally, coercivity showed a ∼3- to 4-fold decrease in the Δ*mamP* strain, and was close to zero in all Δ*mamT* mutant samples.

**Table 5.**
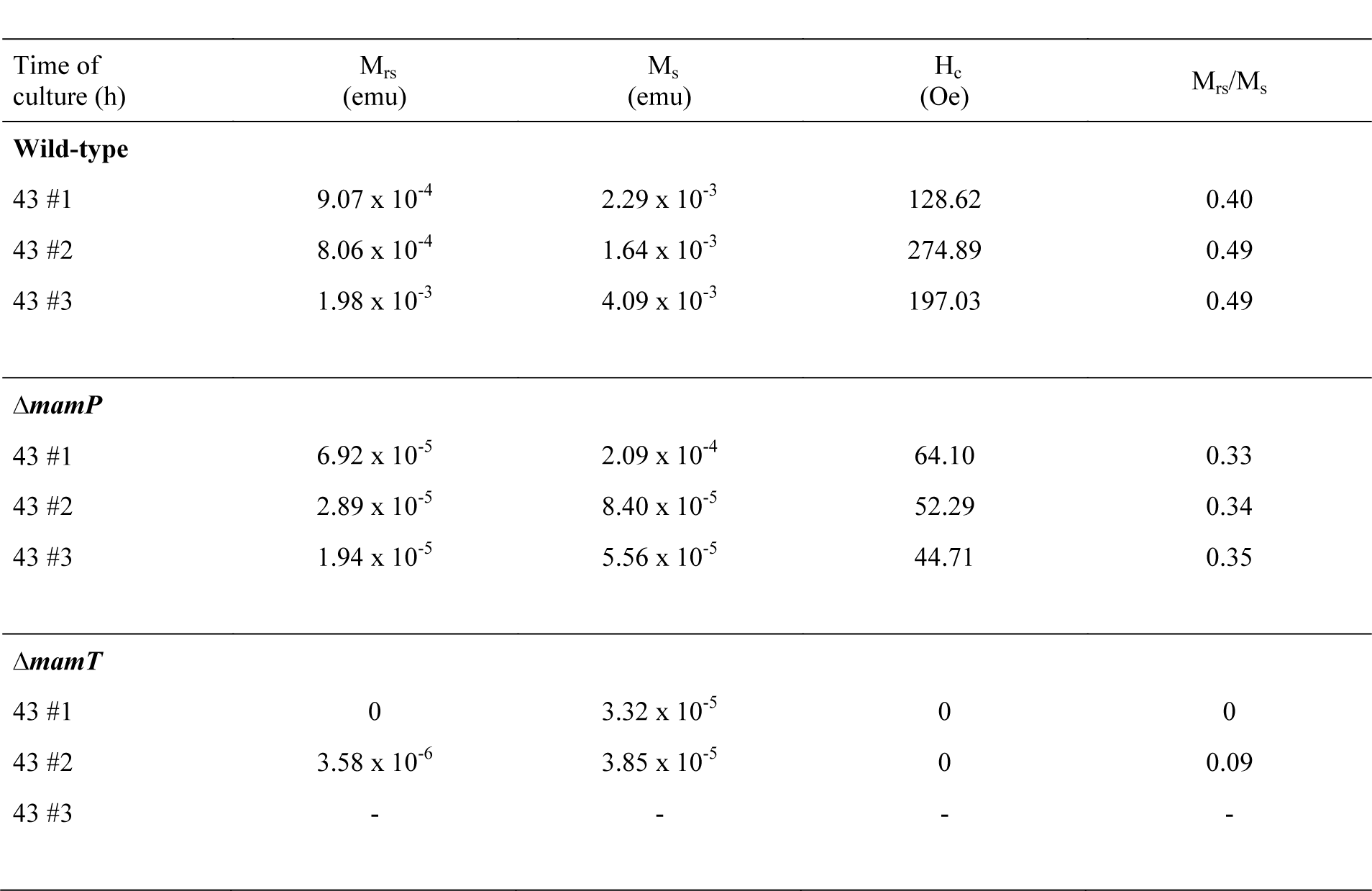
Remanent magnetization (M_rs_), saturation magnetization (M_s_), coercivity (H_c_) and M_rs_/M_s_ ratios of whole mutant cells recovered after cultures. Additional wild-type cultures were used as a control condition.

### Iron distribution in AMB-1 populations

Having determined the mass of iron in the different bacterial pools (*i.e.* magnetite and the rest of the cell), we finally wanted to quantify the iron distribution in AMB-1. The mass of iron in the lysate (mass_lysate_, *i.e.* the fraction distinct from magnetite) was calculated from:

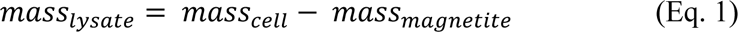

where mass_cell_ and mass_magnetite_ are the mass of iron contained in whole AMB-1 cells and in magnetite, respectively. mass_cell_ was calculated from chemical assays, and mass_magnetite_ was calculated from magnetic characterizations. The fraction of the total cellular iron contained in magnetite (F_magnetite_, in %) was calculated from:

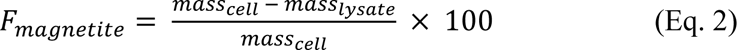

where mass_lysate_ was calculated from Eq. 1.

Using results presented in Figs. 3A and 3B, we calculated the fraction of the total cellular iron contained in magnetite (F_magnetite_ in Eq. 2) (Fig. 6). The mass of magnetite produced by wild-type AMB-1 cells cultivated at 30 and 150 μM was similar after 26 hours of growth, but iron uptake was 10 times higher under high-iron conditions. Therefore, magnetite corresponded to 11.2 ± 6% of the total cellular iron at this time point under low-iron conditions, but only 0.9 ± 0.3% of the cellular iron under high-iron conditions. In bacteria cultivated with 30 μM of iron, the fraction of cellular iron in magnetite increased to 21.9 ± 4 and 26 ± 3% after 46 and 69 hours of culture, respectively. In the 150 μM iron experimental conditions, it increased up to 14.5 ± 3 and 24.3 ± 5% after 46 and 69 hours of culture, respectively (Fig. 6). We note that all cells from every sample observed under the electron microscope contained magnetite crystals.

**FIG 6.**
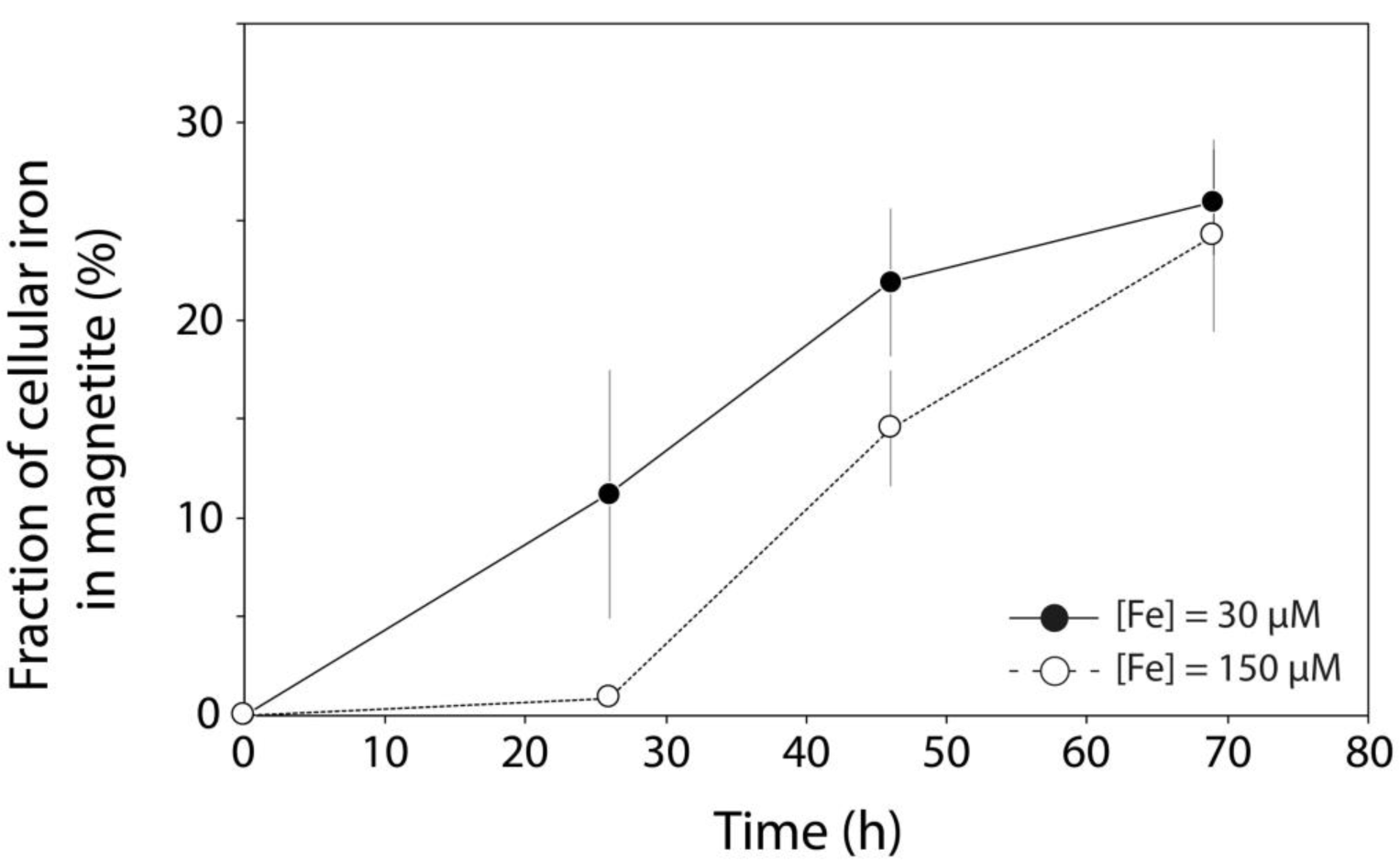
Relative fraction of the total cellular iron contained in magnetite. Each point corresponds to the mean value of three replicates ± 1SD. Black circles and open symbols refer to cultures with an initial iron concentration of 30 and 150 µM, respectively.

Therefore, our data cannot be explained by bacteria that accumulate iron without producing magnetite crystals. We manipulated and stored magnetite under anoxic conditions ([O_2_] < 1ppm) to prevent its oxidation into maghemite [γ-Fe(III)_2_O_3_] (Gallagher et al, 1968; Freer and O’Reilly, 1980; Rebodos and Vikesland, 2010). The saturation magnetization of maghemite is 60-80 emu per gram (Cornell and Schwertmann, 2004). Even if all magnetite became fully oxidized, the mass of iron precipitated as crystals in wild-type AMB-1 would be at most 40% higher as in Fig. 3B, and the mineral fraction of AMB-1 would represent no more than ∼50-60% of the total cellular iron. In this case, our data would still support a significant pool of iron accumulating outside of magnetosome crystals. Finally, we ensured that all iron fractions in AMB-1 cultures were recovered (see materials and methods). Thus, a loss of iron during sample extraction and preparation cannot explain our results. Our data demonstrate that magnetite does not represent the major iron reservoir in AMB-1 under our experimental conditions.

To further demonstrate that iron accumulates outside of magnetite, we used additional wild-type AMB-1 cultures to assess the mean mass of magnetite per AMB-1 cell. Cell counting under a light microscope using a hemocytometer indicated an almost identical total number of cells for the two replicates: 8.48 ± 2.21 x 10^10^ and 8.39 ± 2.26 x 10^10^. The total mass of magnetite produced in these cultures and calculated from magnetic measurements was 0.025 and 0.024 mg. This yields a mean mass of iron contained in magnetite per cell of 2.11 x 10^-7^ and 2.10 x 10^-7^ ng for the two replicates, which corresponds to ∼21 % of the bulk mass of iron measured in AMB-1 cells (Amor *et al*, 2019).

Additional wild-type cultures used as controls for experiments with the mutant strains showed similar results, with ∼31 ± 8 % of the bulk cellular iron contained in magnetite. Magnetite in Δ*mamP* bacteria represented only 13 ± 12 % of the total cellular iron. All Δ*mamP* cells also produced magnetite under our experimental condition. Finally, the fraction of iron contained in Δ*mamT* magnetite could not be determined because of inconclusive data on iron incorporation into these mutant bacteria (see above).

### Subcellular localization and speciation of iron in AMB-1

Iron distribution assessments demonstrated that AMB-1 cells contain a large pool of iron, distinct from magnetite. However, they did not provide physical and chemical information about this additional pool. To determine the localization and speciation of iron in AMB-1, we used the fluorescence FRET Iron Probe 1 (FIP-1), an activity-based probe that allows the detection of labile Fe(II) (Aron et al, 2016). FIP-1 is made of a green (fluorescein) and a red (cyanine) fluorophores linked by an Fe(II)-cleavable endoperoxide. In the native FIP-1 state, the fluorescence energy of the excited fluorescein is transferred to the cyanine through a Fluorescence Resonance Energy Transfer (FRET) mechanism. In that case, only a red fluorescence signal can be observed. Upon reaction with labile Fe(II), the linker between the two fluorophores gets cleaved and a green fluorescence signal can be detected (Aron *et al*, 2016). To further constrain the speciation and subcellular localization of iron distinct from magnetite in AMB-1, wild-type, Δ*mamP* and Δ*mamT* cells were incubated with the FIP-1 probe and imaged via structured illumination microscopy. A mutant strain (ΔMAI) unable to form magnetosomes was used as a negative control (see supplementary materials).

A red fluorescence signal was observed in all samples, indicating the uptake of FIP-1 (Figs. 7, S4 and S5). A very weak green signal was observed in ΔMAI bacteria (Fig. S5), suggesting a lower labile iron concentration in these mutant cells. This observation is in good agreement with quantification of bulk iron in wild-type and ΔMAI bacteria (Amor *et al*, 2019). Both red and green fluorescence patterns showed intracellular heterogeneities, demonstrating that FIP-1 has been internalized into AMB-1. In wild-type bacteria, the green fluorescence signal was diffuse in the cytoplasm, although unstained spaces corresponding to PHB (Poly-β-hydroxybutyrate, a carbon storage molecule) granules can be observed (Figs. 5 and S4). Green fluorescence signal also accumulated at the poles of the cell (Fig. S4). Such accumulation can be observed in dividing cells at the septum location (Fig. S6). Most of wild-type cells also showed a green fluorescence associated with the magnetosome chains (Fig. 7). AMB-1 produces fragmented chains of magnetite, with magnetosome vesicles spreading along the cell’s long axis from pole to pole (Komeili *et al*, 2006; Arakaki *et al*, 2016). Unlike other *Magnetospirillum* strains such as MSR-1, apparent gaps between magnetite crystals can be observed from electron microscopy in AMB-1 (Fig. 1). These gaps correspond to empty magnetosome vesicles, containing no magnetite nanoparticles (Komeili *et al*, 2006; Arakaki *et al*, 2016). In our observations, the green fluorescence signals formed fragmented lines (*i.e.* similar to magnetite crystals) (Figs. 7 and S7). In some rare cases, the fluorescence lines extended almost from poles to poles (*i.e.* similar to magnetosome vesicles). As mentioned above, all bacteria observed with electron microscopy contained magnetite nanoparticles. Therefore, fluorescence patterns showing continuous lines cannot indicate empty vesicles in cells making no magnetite. Δ*mamP* showed all of the fluorescence features that have been observed in the wild-type strain (Figs. 7 and S8), whereas chains of magnetosomes could not be detected in Δ*mamT* AMB-1 using FIP-1 (Figs. 7 and S9).

**FIG 7.**
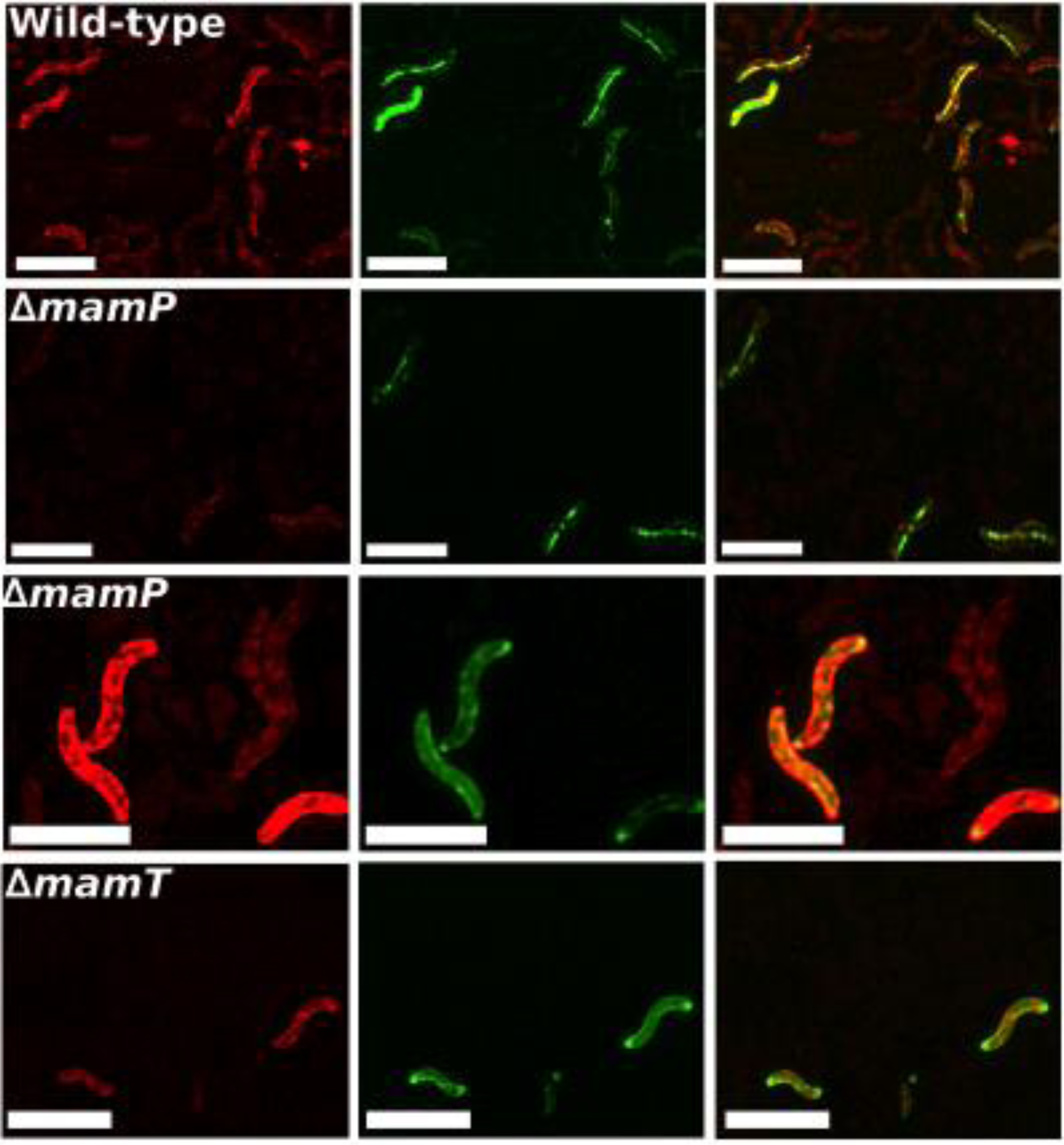
Red (left panels), green (center panels) and merged (right panels) fluorescence images of wild-type, Δ*mamP* and Δ*mamT* AMB-1 incubated with FIP-1 for 180 min. The two Δ*mamP* panels show the two fluorescence patterns (diffuse and located to the magnetosome chains, as in wild-type) observed in the populations. Scale bars = 4 microns. Additional pictures are available in the supplementary materials.

## Discussion

Mass balance experiments identified a large amount of iron distinct from magnetite in AMB-1, representing ∼75 % of the bulk cellular iron in our experimental conditions. These results suggest a more complex picture for iron cycling and homeostasis in MTB than previously thought, as intracellular iron needs to be handled by the cell to prevent toxic effects.

### Iron incorporation and distribution in AMB-1

Monitoring iron concentration and oxidation state in AMB-1 growth medium demonstrated that initial Fe(III) became progressively reduced into Fe(II) (Fig. 4). Accumulation of Fe(II) could result from active reduction by AMB-1, or illustrate respiration reactions depleting oxygen in AMB-1 medium. Our experimental setup cannot rule out one of the two possibilities, but we note that iron isotopes identified Fe(III) reduction within AMB-1 cells, and subsequent diffusion of intracellular Fe(II) to the growth medium (Amor *et al*, 2018).

Iron incorporation into wild-type AMB-1 was higher under high-iron conditions. When normalized to optical density, which is proportional to the concentration of cells in culture, iron uptake by AMB-1 after 26 h of culture was ∼10-fold higher in the 150 μM experimental condition, compared to the 30 μM condition (Fig. 3). However, the mass of magnetite was similar in the two culture conditions (Fig. 3), indicating that the limiting step for biomineralization corresponds to magnetite precipitation and maturation, rather than iron uptake into the cell. Mass balance estimations were consistent in all wild-type cultures, and indicated that ∼25 to ∼30 % of the bulk cellular iron was contained in magnetite after 69h of growth. The mean mass of iron contained in magnetite per cell (∼0.21 x 10^-6^ ng), estimated from cell counting and magnetic quantification using a VSM, corresponds to 21 % of the bulk iron content in AMB-1 determined by single-cell mass spectrometry analyses under the same experimental conditions (Amor *et al*, 2019). The results are almost identical to the mass balance estimations, and show that most of iron is contained in reservoir(s) distinct from magnetite. Moreover, the combination of electron microscopy and mass spectrometry measurements for quantification of iron content in AMB-1 evidenced a delay in magnetite formation as iron was incorporated into bacteria (Amor *et al*, 2019). This observation further supports accumulation of intracellular iron outside of magnetite.

The limited iron incorporation into Δ*mamP* AMB-1 suggests that magnetite biomineralization regulates iron assimilation. Whether this regulation corresponds to a direct or indirect mechanism remains unclear. A likely hypothesis could be that the iron accumulation capacity of the cell’s fraction distinct from magnetite is limited. Once bacteria are fully loaded with iron, its sequestration into magnetite would be required for further assimilation. Such model would imply a two-step process for magnetite biomineralization, in which iron is first stored in the non-crystalline fraction of the cell, and then precipitated as magnetite. This is in good agreement with what has been proposed for iron cycling in AMB-1 and MSR-1 (Baumgartner *et al*, 2013; Fdez-Gubieda *et al*, 2013; Amor *et al*, 2018, 2019). The lack of MamP in the mutant strain could hamper iron precipitation in magnetosomes, and thus indirectly prevent further iron assimilation.

If 75 % of intracellular iron in AMB-1 is not stored in magnetite, the pool distinct from magnetite should represent ∼0.75 x 10^-6^ ng of iron per cell (Amor *et al*, 2019). This mass is estimated to be 10 to 100-fold higher than the mass of iron in *Escherichia coli* cells (Andrews *et al*, 2003). The iron content in the ΔMAI AMB-1 strain, unable to form magnetosomes, was also estimated to be 5- to 10-fold higher than *E. coli* cells (Amor *et al*, 2019). Excess of free iron in the intracellular medium is toxic for cells (Andrews *et al*, 2003), which suggests efficient iron storage and detoxifying pathways in AMB-1. They could include ferritins, bacterioferritins and Dps proteins (Andrews *et al*, 2003; Uebe *et al*, 2010, 2019). Dps and bacterioferritins have recently been shown to protect MSR-1 from oxidative stress (Uebe *et al*, 2019), and phases corresponding to ferritin-like structures have been evidenced in AMB-1, MSR-1, and MS-1 strains using spectroscopic methodologies (Frankel et al, 1983; Faivre *et al*, 2007; Uebe *et al*, 2010; Baumgartner *et al*, 2013; Fdez-Gubieda *et al*, 2013; Uebe *et al*, 2019). Further iron toxicity assays in MTB and mutant strains lacking some of these iron-storing proteins will help to better understand the capacity of MTB to tolerate such high intracellular iron concentrations.

### Discrepancy with previous work

Our results clearly showed discrepancy with previous characterizations of iron species in the two magnetotactic strains AMB-1 and MSR-1 using X-ray absorption methodologies (Baumgartner *et al*, 2013; Fdez-Gubieda *et al*, 2013). In these two studies, time-course experiments were carried out in which AMB-1 or MSR-1 cells were cultivated without iron. When reaching saturation of cell density, iron was added to the growth medium to trigger magnetite biomineralization. Phases likely corresponding to ferrihydrite were first observed in both strains. When biomineralization was complete, magnetite was the sole iron carrier observed in bacteria. Since additional iron was detected by mass spectrometry and the protocol described in the present research, such discrepancy could suggest that X-ray absorption is not suitable for detection of iron that is not contained in magnetite or ferrihydrite. However, the important fraction of iron we identified in the iron pool distinct from magnetite rather suggests that the discrepancy arises from different experimental protocols. Moreover, recent work on MSR-1 also suggested that iron can be contained outside of magnetite in this strain (Berny *et al*, 2020). Iron-starving conditions can impact the iron cycling and homeostasis in MTB, as low-iron conditions have been shown to induce overexpression of iron acquisition systems in AMB-1 and MSR-1 (Suzuki *et al*, 2006; Wang *et al,* 2017). They might optimize the transfer of incorporated iron to magnetosomes for magnetite precipitation. Further X-ray absorption analyses with iron-starved bacteria and cells grown under standard conditions (*i.e.* as in the present work) will be needed to confirm this hypothesis.

### The magnetic properties of AMB-1 cultures illustrate defects in magnetite biomineralization

In addition to the mass of magnetite produced in AMB-1 cultures, magnetic characterizations of bacterial samples provided important insights on the nanoparticle size and organization. AMB-1 produces stable single-domain magnetite nanoparticles (Li *et al*, 2012). The M_rs_ / M_s_ ratios corresponding to wild-type, mature AMB-1 magnetite typically range between 0.43 and 0.50 (Dunlop, 2020; Li *et al*, 2012). Smaller magnetite particles with crystal dimensions below 30 nm (for a width / length ratio of 0.2 or higher) are not magnetically stable at room-temperature, and fall within the superparamagnetic domain (Muxworthy and Williams, 2009). Their remanence magnetization is thus 0 at room-temperature, but their saturation magnetization remains unchanged for a given mass of magnetite. Very small superparamagnetic particles (<10 nm in length) would not reach complete saturation under the maximum external field we used (4 000 Oe) at room-temperature (Zuoquan et al, 2014). These particles represent less than 5% of the magnetite crystals observed under electron microscopy (Fig. S2). The underestimation of the saturation magnetization of these nanoparticles would be around 20 % (Zuoquan et al, 2014), meaning that the underestimation of the mass of magnetite would be below 1% and thus negligible. A mixing of stable single-domain and superparamagnetic particles would lead to lower M_rs_ / M_s_ ratios (Dunlop, 2002). M_rs_ / M_s_ ratios of ∼0.38 observed after 26h of growth in wild-type cultures could thus reflect a mixing of mature and newly formed particles. Even though iron uptake was ∼10-fold higher under high-iron conditions (Fig. 3), M_rs_ / M_s_ ratios in the two iron conditions were identical regardless of the initial iron concentration in the growth medium. As mentioned above, this suggests magnetite precipitation and growth as the limiting step for biomineralization in AMB-1. For longer culture times, M_rs_ / M_s_ ratios in wild-type AMB-1 were consistent with values ranging between 0.46 and 0.49 typical of AMB-1 magnetite (Li *et al*, 2012). Δ*mamP* and Δ*mamT* AMB-1 also showed decreased M_rs_ / M_s_ ratios compared to the expected ∼0.45 value. M_rs_ / M_s_ ratios in Δ*mamP* cultures of 0.34 are consistent with a mixing of stable single-domain magnetite and smaller superparamagnetic particles as confirmed by electron microscopy observations (Fig. 1). As mentioned above, very small superparamagnetic particles would not reach saturation at room-temperature, leading to an underestimation of the mass of magnetite of ∼20%. In that case, our results would still support a large pool of iron distinct from magnetite in the Δ*mamP* strain. Contrastingly, Δ*mamT* showed M_rs_ and M_rs_ / M_s_ ratios of zero, which are both consistent with superparamagnetic particles produced in this strain. Electron microscopy demonstrated that Δ*mamT* bacteria produced ∼10 to 20-nm long nanoparticles, in good agreement with the magnetic analyses (Muxworthy and Williams, 2009).

Finally, coercivity in wild-type cultures increased with time in both iron conditions. Coercivity in stable single-domain particles such as those produced by MTB (Li et al, 2012) depends on the particle size and shape, as well as the magnetocrystalline anisotropy which is unlikely to change. Consistent H_c_ values between the two iron conditions after 26h of culture suggest similar size and shape for magnetite crystals, in good agreement with iron uptake patterns, remanent magnetization, and saturation magnetization (see above). For longer times of culture, AMB-1 cultivated with iron at 150 μM showed higher coercivity. Since M_rs_ is unchanged between these two cultures, this requires that the particles are larger or a different shape under high-iron conditions. To test this hypothesis, we measured the magnetite length and width distributions in both experimental conditions. Results are given in Figs. S2 and S3, and show that the shape (length/width ratio) is almost identical in both experimental conditions. However, bigger magnetite crystals were produced by AMB-1 cultivated with iron at 150 μM (mean length of 38.46 nm) compared to the 30 μM condition (mean length of 32.03 nm), suggesting that increased coercivity results from bigger particles under high-iron conditions. Lower coercivities in the mutant strains can also be explained by the presence of small superparamagnetic particles.

### Localizing iron in AMB-1 cells

Two green fluorescence patterns were observed in AMB-1 cultures: a diffuse signal across the cell, and a signal that concentrated on the magnetosome chain. The SIM fluorescence microscope did not allow image acquisition of standard optical observation using white light, thus preventing estimation of the fraction of the total cells showing a fluorescence signal. Observation of magnetosome chains using FIP-1 shows that Fe(II) is addressed to magnetosomes during biomineralization. There is a possibility that FIP-1 indicates poorly crystalline Fe(II) at the magnetite surface, but our observations are best explained by Fe(II) being contained either in magnetite-containing or magnetite-free magnetosome vesicles (see below). It is unclear whether this Fe(II) would be contained inside magnetosomes, within the magnetosome membrane, or at the magnetosome surface. From high-resolution electron microscope analyses, Werckmann and collaborators proposed that iron could accumulate in the magnetosome membrane before its precipitation as magnetite (Werckmann *et al*, 2017). Our observations are in line with these results, and indicate that iron in the magnetosome membrane would at least be composed of Fe(II) species. Genes encoding for Fe(II) transporters have been found in the magnetosome gene island, and could transport Fe(II) across the magnetosome membrane for magnetite formation (Suzuki *et al*, 2006; Rong *et al*, 2008, 2012). Identical fluorescence patterns were observed in Δ*mamP* AMB-1, but not in the Δ*mamT* strain. The chain-like localization pattern in Δ*mamP* suggests that FIP-1 does not bind to magnetite, since this mutant strain produces only a few crystals per cell. It also indicates that Fe(II) is delivered to magnetosomes in the AMB-1 cells lacking MamP, and suggests that magnetite-free magnetosomes can be stained by FIP-1.

Another notable observation is an accumulation of fluorescence at the poles of AMB-1 cells showing a diffuse green signal. Whether such accumulation illustrates true biological mechanisms (flagellar apparatus, chemotaxis receptors, nitrate reductase complex, siderosomes, cell division) remains speculative, and additional work will be necessary to determine the significance of these observations (Müller et al, 2014; Popp *et al*, 2014; Alberge *et al*, 2015; Cunrath *et al*, 2015; Gasser *et al*, 2015).

Lastly, it is important to note that FIP-1 does not react with Fe(II) bound tightly to proteins as well as Fe(III) (Aron et al, 2016). It is likely that additional iron species distinct from magnetite are contained in AMB-1 cells, which include iron associated with heme domains (Siponen *et al*, 2013; Jones *et al*, 2015) or iron contained in storage proteins such as ferritins (Faivre et al, 2007; Uebe *et al*, 2019).

### Implications for Earth sciences

It has been hypothesized that MTB deplete their environment in bioavailable iron by sequestering dissolved species into magnetite (Lin *et al*, 2014). Once the cell dies, MTB magnetite crystals can be trapped into sedimentary rocks, which effectively removes iron from the dissolved pool (Kopp and Kirschvink, 2008, Larrasoaña *et al*, 2014). MTB could thus prevent other living organisms from accessing an available source of iron. Some parameters are missing to accurately quantify the impact of MTB on the iron biogeochemical cycle. One of them is the speciation of iron in MTB, which controls its solubility. Our data demonstrated that most of iron in MTB exists as soluble species (*i.e.* Fe(II) and soluble Fe(III)-organic compounds), rather than magnetite. Iron sequestration in environmental MTB might thus be more limited than previously proposed (Amor *et al*, 2019). However, the discrepancy between the present work and the former X-ray absorption characterizations of iron in MTB (Baumgartner *et al*, 2013; Fdez-Gubieda *et al*, 2013) raises questions about the environmental significance of our findings. Environmental MTB populations could experience varying iron conditions, and transition from iron-starving to iron-rich conditions similar to X-ray absorption experiments (Baumgartner *et al*, 2013; Fdez-Gubieda *et al*, 2013). In this case, most of intracellular iron could be contained in magnetite, with a limited soluble iron pool. Additional work constraining iron speciation in MTB that experience transitioning iron conditions, as well as in bacterial populations sampled from the environment, will be useful to further address the impact of MTB on the iron biogeochemical cycle.

## Materials and Methods

### Cultivation of wild-type and mutant AMB-1 strains

*Magnetospirillum magneticum* AMB-1 (ATCC 700264) was cultivated in 200 mL-bottles. The detailed composition of AMB-1 growth medium is given by Komeili and co-workers (Komeili *et al*, 2004). The sole iron source provided to AMB-1 cultures corresponded to Fe(III)-citrate, which was added to the growth media from an Fe(III)-citrate solution prepared by mixing Fe(III)Cl_3_ (6 mM) and citric acid (12 mM) powders (Sigma-Aldrich) in Milli-Q water. The pH of the Fe(III)-citrate solution was set at 6.9 (*i.e.* same as AMB-1 growth medium) using NaOH. The initial Fe(III) concentration in AMB-1 growth medium was either 30 (*i.e.* standard concentration used in the ATCC medium) or 150 (*i.e.* the concentration used for isotope experiments) μM. The concentration of citrate and volume of cultures were kept constant in all experiments by adding an iron-free citrate solution (12 mM, pH 6.9) to AMB-1 cultures under low-iron conditions (30 μM). AMB-1 was cultivated in a glove box with controlled atmosphere (90% N_2_, 10% O_2_) at 30°C for three days. Each day, one bottle for each experimental condition was recovered for chemical and magnetic characterizations (see below). All measurements were carried out in triplicates (total of 18 bottles: two iron conditions, three time points, three replicates for each condition).

Two AMB-1 mutant strains were selected for additional experiments: the *ΔmamP* and Δ*mamT* strains lacking the genes encoding for the MamP and MamT proteins, respectively (Murat *et al*, 2010). MamP and MamT are magnetochrome proteins, a class of c-type cytochromes specific to MTB, which can bind iron via their heme domains (Siponen *et al*, 2013). Magnetochromes have been proposed to regulate the iron oxidation state in MTB (Siponen *et al*, 2013). The two mutant strains show biomineralization defects, which enable us to investigate the link between magnetite formation and iron uptake. The *ΔmamP* strain produces only a few crystals per cell resembling those produced by wild-type AMB-1, as well as a few additional small crystals (Fig. 1A-C). Δ*mamT* bacteria synthesize many small, elongated crystals (Fig.1D). The *ΔmamP* and *ΔmamT* strains have already been produced by our group (Murat *et al*, 2010). In AMB-1, *mamT* gene is located in the *mamAB* gene clusters of the MAI (termed R5 region in our previous work) downstream of three genes *mamQ*, *mamR* and *mamB* (Murat *et al*, 2010; Uebe and Schüler, 2016). These three genes are perfectly duplicated in the R9 region of the MAI, downstream of *mamT*. To avoid recombination between regions R5 and R9, the region R9 was deleted from *ΔmamT*. Therefore, bacteria used in this study correspond to the *ΔmamTΔR9* strain, and are referred to as Δ*mamT*. We ensured that *ΔmamT* and *ΔmamTΔR9* strains show similar biomineralization defects and both can be complimented with *mamT* expressed from a plasmid (Jones *et al*, 2015). Because the mutant strains produce less magnetite than wild-type AMB-1, as shown by the electron microscope observations (Fig. 1), we cultivated them with Fe(III)-citrate at 150 μM to measure iron uptake more accurately. The highest iron uptake by wild-type AMB-1 was observed at ∼45 h of culture (see results). Accordingly, *ΔmamP* and *ΔmamT* strains were cultivated for ∼45 hours in triplicates in 200-mL bottles (total of 9 bottles: 3 replicates for *ΔmamP* and *ΔmamT* and 3 replicates for wild-type bacteria used as a control).

### Transmission electron microscopy

Bacteria were deposited on copper grids coated with a Formvar film, and observed with a transmission electron microscope (FEI Tecnai 12) operating at 120 kV. From electron microscopy observations, the length of magnetite nanoparticles produced by wild-type AMB-1 cultivated for three days with Fe(III) at either 30 or 150 μM was measured using the ImageJ software.

### Chemical measurements

Bacterial iron uptake was quantified by measuring iron concentration in AMB-1 cultures at initial (immediately after inoculation) and final (at the end of the bacterial culture) stages using the ferrozine assay (Hunter *et al*, 2013). Ferrozine forms a purple-colored complex with Fe(II), which can be determined spectrophotometrically. Total iron is then determined by total reduction of iron in the sample with hydroxylamine hydrochloride and subsequent reaction with ferrozine. Concentration of Fe(III) is calculated as the difference of total iron and Fe(II). For each condition, pH and optical density at 400 nm (OD_400nm_) were measured. Then, 1 mL of culture was sampled and filtered (0.22-μm pore size; Acrodisc syringe filters, polyethersulfone) at the initial and final stages. The Fe(II) and total iron concentrations were measured using the ferrozine assay. The mass of iron taken up by AMB-1 was calculated from iron depletion in each culture.

To demonstrate the reliability of the ferrozine assay for measuring iron depletion in AMB-1 cultures, we also prepared sterile growth media provided with Fe(III)-citrate at 30 or 150 μM in 200-mL bottles. One mL of growth media was sampled and filtered (0.22-μm pore size; Acrodisc syringe filters, polyethersulfone) after iron addition. Iron concentration and speciation was measured with the ferrozine assay as described above. Sterile bottles were incubated for 1, 2 or 3 days at 30°C in the glove box (90% N_2_, 10% O_2_). At the end of experiment, 1 mL of growth media was recovered, filtered and the iron concentration and speciation were measured using the ferrozine assay. Three replicates per condition were prepared (18 samples total, as for wild-type bacterial cultures).

### Magnetic characterizations

After chemical analyses, whole growth media were recovered and centrifuged (8,000 rpm, 10 min). Supernatants were discarded, and bacterial pellets corresponding to the entire bacterial populations were dried in an anoxic chamber (98% N_2_, 2% H_2_, O_2_ < 1 ppm) at room temperature to prevent magnetite oxidation. We have already demonstrated that no significant fraction of iron is adsorbed on AMB-1 cell surfaces (Amor *et al*, 2018). Virtually all iron contained in bacterial pellets thus corresponds to intracellular iron. Once dried, whole bacterial pellets were transferred into sample holders inside the anoxic chamber for subsequent magnetic characterizations. Samples were kept in anoxic conditions until magnetic analyses were performed. Hysteresis loops of magnetization versus applied magnetic field were measured using a Vibrating-Sample Magnetometer (LakeShore VSM 7410) at room temperature. An integration time of 10 s per point was used.

### Iron mass balance

To demonstrate the validity of our protocol and the accuracy of our measurements, we ensured that all iron fractions were recovered and that no iron was lost during sample extraction and preparation. Additional wild-type AMB-1 cultures were carried out in 200-mL bottles for three days. One mL of growth medium was sampled and filtered before and after cultures, and iron concentration was measured using the ferrozine assay as described above. Cells were recovered by centrifugation (8,000 rpm, 10 min). The supernatant was discarded and bacterial pellets were suspended in 100 μL of phosphate buffer (PBS). Cells were washed three times in PBS and stored for subsequent measurements of total cellular iron mass (m_cell_) using single-cell – inductively coupled plasma - mass spectrometry following a protocol previously published (Amor *et al*, 2019). Before mass-spectrometry measurements, PBS solution containing the bacteria was filtered to measure the potential mass of iron that leaked outside of the cells using high-resolution – inductively coupled plasma - mass spectrometry (m_leaked_) (Amor *et al*, 2019). Iron recovery was assessed from the following mass balance equation:

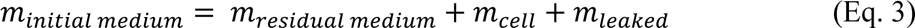

where m_initial medium_, and m_residual medium_ are the mass of iron in the growth medium before and after AMB-1 cultures, respectively. m_leaked_ represented ∼0.5 % of m_cell_ or less (Table 1). Mass balance estimations showed that iron recovery during sample preparation was ranging between 96 and 100% (Table S1), demonstrating the validity of our protocol. Therefore, a loss of iron pools such as magnetite could not explain our results.

### Cell counting

To further demonstrate that wild-type AMB-1 incorporates more iron than needed to make its magnetite crystals, we chose an alternative approach to estimate the mean mass of magnetite per cell and to compare these results with available data on single-cell bulk iron content in AMB-1. Additional wild-type AMB-1 cultures (two replicates) were grown with Fe(III)-citrate at 150 μM. For each culture, the entire AMB-1 population was recovered with centrifugation, and the total mass of magnetite in a given population was determined from magnetic measurements as described above. The number of cells in the same populations was then calculated from direct cell counting using a hemocytometer under a light microscope.

Fifty counts were done for each AMB-1 culture. Finally, the mean mass of iron per cell was calculated from the total number of cells and the total mass of magnetite in each culture.

### Detection and mapping of labile Fe(II) in AMB-1 using the FIP-1 fluorescent probe

The sensing mechanism for FIP-1 is inspired by antimalarial natural products and related therapeutics (Borstnik et al, 2002; Tang et al, 2005; Creek et al, 2007; Spangler et al, 2016). This reagent has been developed for use in mammalian cells and expanded in bioluminescent versions for mouse imaging (Aaron *et al*, 2017). We adapted the use of FIP-1 (Aron *et al*, 2016) for detection of Fe(II) in AMB-1. Wild-type and mutant AMB-1 strains were cultivated in 10 mL glass tubes until end of exponential phase / beginning of the stationary phase. Ten mL of growth medium were centrifuged, the supernatant was discarded, and cells were resuspended in 500 μL of PBS buffer. To ensure that all iron from the growth medium is removed, cells were centrifuged and washed in fresh PBS buffer three times. Finally, the three bacterial strains were mixed with a PBS solution containing EDTA (5mM, pH 6.9) for 10 min, centrifuged, and suspended in the FIP-1 solution (*i.e* FIP-1 at 1 mM in Hank’s Balance Salt Solution) for 180 min at 30°C in the glove box (90% N_2_, 10% O_2_). All bacterial samples were observed by Structured Illumination Microscopy with a Carl Zeiss Elyra PS.1 Super Resolution fluorescence microscope, using red (excitation wavelength of 561 nm, emission wavelength of 570-620 nm) and green (excitation wavelength of 488 nm, emission wavelength of 495-550 nm) laser lines for the detection of the native and cleaved probe, respectively. Images were processed with the ImageJ software.

## ACKNOWLEDGMENTS

AK and MA are supported by grants through the National Science Foundation (1504681) and National Institute of Health (R01GM084122 and R35GM127114). AC, CS and FH were supported by the Director, office of Science, Office of Basic Energy Sciences, Materials Sciences and Engineering Division, of the U.S. Department of Energy under Contract No. DE-AC02-05-CH11231 within the Nonequilibrium Magnetic Materials Program (KC2204). CJC and ATA were supported by a grant from the National Institute of Health (R01GM079465). CJC is a CIFAR Senior Fellow. ATA thanks the NSF for a graduate fellowship and was partially supported by an NIH Chemical Biology Interface Training Grant (T32 GM066698). We thank Mickaël Tharaud for assistance with mass spectrometry analyses. Part of this work was supported by IPGP multidisciplinary program PARI and by Region Ile-de-France SESAME Grant no. 12015908.

## References

Aharoni A. (2000) Introduction to the theory of ferromagnetism. Oxford University Press, Oxford.

Alberge F., Espinosa L., Seduk F., Sylvi L., Toci R., Walburger A., Magalon A. (2015) Dynamic subcellular localization of a respiratory complex controls bacterial respiration. eLife 4, #e05357 doi: 10.7554/eLife.05357.

Amor M., Tharaud M., Gélabert A., Komeili A. (2019) Single-cell determination of iron content in magnetotactic bacteria: implications for the iron biogeochemical cycle. Environ. Microbiol. In press. https://doi.org/10.1111/1462-2920.14708.

Amor M., Busigny V., Louvat P., Tharaud M., Gélabert A., Cartigny P., Carlut J., Isambert A., Durand-Dubief M., Ona-Nguema G., Alphandéry E., Chebbi I., Guyot F. (2018) Iron uptake and magnetite biomineralization in the magnetotactic bacterium *Magnetospirillum magneticum* strain AMB-1: an iron isotope study. Geochim. Cosmochim. Acta 232, 225–243.

Amor M., Busigny V., Louvat P., Gélabert A., Cartigny P., Durand-Dubief M., Ona-Nguema G., Alphandéry E., Chebbi I., Guyot F. (2016) Mass-dependent and -independent signature of Fe isotopes in magnetotactic bacteria. Science 352, 705–708.

Andrews S. C., Robinson A. K., Rodríguez-Quiñones F. (2003) Bacterial iron homeostasis. FEMS Microbiol. Rev. 27, 215–237.

Arakaki A., Kikuchi D., Tanaka M., Yamagishi A., Yoda T., Matsunaga T. (2016) Comparative subcellular localization analysis of magnetosome proteins reveals a unique localization behavior of Mms6 protein onto magnetite crystals. J. Bacteriol. 198, 2794–2802.

Aron A. T., Loeehr M. O., Bogena J., Chang C. J. (2016) An endoperoxide reactivity-based FRET Probe for ratiometric fluorescence imaging of labile iron pools in living cells. J. Am. Chem. Soc. 138, 14338–14346.

Aron A. T., Heffern M. C., Lonergan Z. R., Vander Wal M. N., Blank B. R., Spangler B., Zhang Y., Park H. M., Stahl A., Renslo A. R., Skaar E. P., Chang C. J. (2017) *In vivo* bioluminescence imaging of labile iron accumulation in a murine model of *Acinobacter baumannii* infection. Proc. Natl. Acad. Sci. U. S. A. 114, 12669–12674.

Baumgartner J., Morin G., Menguy N., Perez Gonzalez T., Widdrat M., Cosmidis J., Faivre D. (2013) Magnetotactic bacteria form magnetite from a phosphate-rich ferric hydroxide via nanometric ferric (oxyhydr)oxide intermediates. Proc. Natl. Acad. Sci. U. S. A. 110, 14883–14888.

Berny C., Le Fèvre R., Guyot F., Blondeau K., Guizonne C., Rousseau E., Bayan N., Alphandéry E. (2020) A method for producing highly pure magnetosomes in large quantity for medical applications using *Magnetospirillum gryphiswaldense* MSR-1 magnetotactic bacteria amplified in minimal growth media. Front. Bioeng. Biotechnol. 8, 16. Doi: 10.3389/fbioe.2020.00016.

Borstnik K., Paik I. H., Shapiro T. A., Posner G. H. (2002) Antimalarial chemotherapeutic peroxides: artemisinin, yingzhaosu A and related compounds. Int. J. Parasitol. 32, 1661–1667.

Cornell R. M., Schwertmann U. (2013) The iron oxides: structure, properties, reactions, occurrences and uses, second edition. Wiley-VCH Verlag GmbH & Co. KGaA.

Creek D. J., Charman W. N., Chiu F. C., Prankerd R. J., McCullough K. J., Dong Y., Vennerstrom J. L., Charman S. A. (2007) Iron-mediated degradation kinetics of substituted dispiro-1,2,4-trioxolane antimalarials. J. Pharm. Sci. 96, 2945–2956.

Cunrath O., Gasser V., Hoegy F., Reimmann C., Guillon L., Schalk I. J. (2015) A cell biological view of the siderophore pyochelin iron uptake pathway in *Pseudomonas aeruginosa*. Environ. Microbiol. 17, 171–185.

Dunlop D. J. (2002) Theory and application of the Day plot (M_rs_/M_s_ versus H_cr_/H_c_): 1/ Theoretical curves and tests using titanomagnetite data. J. Geophys. Res. 107, doi: 10.1029/2001JB000486.

Faivre D., Böttger L. H., Matzanke B. F., Schüler D. (2007) Intracellular magnetite biomineralization in bacteria proceeds by a distinct pathway involving membrane-bound ferritin and an iron(II) species. Ang. Chem. Int. Ed. 46, 8495–8499.

Fdez-Gubieda M. L., Muela A., Alonso J., García-Prieto A., Olivi L., Fernández-Pacheco R., Barandiarán J. M. (2013) Magnetite biomineralization in *Magnetospirillum gryphiswaldense*: time-resolved magnetic and structural studies. ACS Nano 7, 3297–3305.

Frankel R. B., Papaefthymiou G. C., Blakemore R. P., O’Brien W. (1983) Fe_3_O_4_ precipitation in magnetotactic bacteria. Biochim. Biophys. Acta. 763, 147–159.

Freer W., O’Reilly R. (1980) The diffusion of Fe^2+^ ions in spinels with relevance to the process of maghemitization. Mineral. Mag. 43, 889–899.

Gallagher K. J., Feitknecht W., Mannweiler U. (1968) Mechanism of oxidation of magnetite to γ-Fe_2_O_3_. Nature 217, 1118–1121.

Gasser V., Guillon L., Cunrath O., Schalk I. J. (2015) Cellular organization of siderophore biosynthesis in Pseudomonas aeruginosa: evidence for siderosomes. J. Inorg. Biochem. 148, 27–34.

Hunter R. C., Asfour F., Dingemans J., Osuna B. L., Samad T., Malfroot A., Cornelis P., Newman D. K. (2013) Ferrous iron is a significant component of bioavailable iron in cystic fibrosis airways. mBio 4, e00557–13.

Jones S. R., Wilson T. D., Brown M. E., Rahn-Lee L., Yu Y., Fredriksen L. L., Ozyamak E., Komeili A., Chang M. C. Y. (2015) Genetic and biochemical investigations of the role of MamP in redox control of iron biomineralization in *Magnetospirillum magneticum*. Proc. Natl. Acad. Sci. U. S. A. 112, 3904–3909.

Komeili A., Li Z., Newman D. K., Jensen G. J. (2006) Magnetosomes are cell membrane invaginations organized by the actin-like protein MamK. Science 31, 242–245.

Komeili A., Vali H., Beveridge T. J., Newmann D. K. (2004) Magnetosome formation are present before magnetite formation, and MamA is required for their activation. Proc. Natl. Acad. Sci. U. S. A. 101, 3839–3844.

Kopp R. E., Kirschvink J. L. (2008) The identification and biogeochemical interpretation of fossil magnetotactic bacteria. Earth-Sci. Rev. 86, 42–61.

Larrasoaña J. C., Liu Q., Hu P., Roberts A. P., Mata P., Civis J., Sierro F. J., Pérez-Asensio J. N. (2014) Paleomagnetic and paleoenvironmental implications of magnetofossil occurences in late Miocene marine sediments from the Guadalquivir basin, SW Spain. Front. Microbiol. 5, #71 doi: 10.3389/fmicb.2014.00071.

Li J., Pan Y., Chen G., Liu Q., Tian L., Lin W. (2009) Magnetite magnetosome and fragmental chain formation of *Magnetospirillum magneticum* AMB-1: transmission electron microscopy and magnetic observations. Geophys. J. Int. 177, 33–42.

Li J., Wu W., Liu Q., Pan Y. (2012) Magnetic anisotropy, magnetostatic interactions and identification of magnetofossils. Geochem. Geophys. Geosyst. 13, #Q10Z51. doi: 10.1029/2012GC004384.

Lin W., Bazylinski D. A., Xiao T., Wu L.-F., Pan Y. (2014) Life with compass: diversity and biogeography of magnetotactic bacteria. Environ. Microbiol. 16, 2646–2658.

McCausland H. C., Komeili A. (2020) Magnetic genes: studying the genetics of biomineralization in magnetotactic bacteria. PLoS Genet. 16, e1008499. Doi: 10.1371/journal.pgen.1008499.

Meister M. (2016) Physical limits to magnetogenetics. eLife 5, e17210. doi: 10.7554/eLife.17210.

Müller F., Raschdorf O., Nudelman H., Messerer M., Katzmann E., Plitzko J. M., Zarivach R., Schüler D. (2013) The FtsZ-like protein FtsZm of *Magnetospirillum gryphiswaldense* likely interacts with its generic homolog and is required for biomineralization under nitrate deprivation. J. Bacteriol. 196, 650–659.

Murat D., Quinlan A., Vali H., Komeili A. (2010) Comprehensive genetic dissection of the magnetosome gene island reveals the step-wise assembly of a prokaryotic organelle. Proc. Natl. Acad. Sci. U. S. A. 107, 5593–5598.

Muxworthy A. R., Williams W. (2009) Critical superparamagnetic/single-domain grain sizes in interacting magnetite particles: implications for magnetosome crystals. J. R. Soc. Interface 6, 1207–1212.

Popp F., Armitage J. P., Schüler D. (2014) Polarity of bacterial magnetotaxis is controlled by aerotaxis through a common sensory pathway. Nat. Commun. 5, #5398 doi: 10.1038/ncomms6398.

Rebodos R. L., Vikesland P. J. (2010) Effects of oxidation on the magnetization of nanoparticulate magnetite. Langmuir 26, 16745–16753.

Rong C., Zhang C., Zhang Y., Qi L., Yang J., Guan G., Li Y., Li J. (2012) FeoB2 functions in magnetosome formation and oxidative stress protection in *Magnetospirillum gryphiswaldense* strain MSR-1. J. Bacteriol. 194, 3972–3976.

Rong C., Huang Y., Zhang W., Jiang W., Li Y., Li J. (2008) Ferrous iron transport protein B gene (*feoB1*) plays an accessory role in magnetosome formation in *Magnetospirillum gryphiswaldense* strain MSR-1. Res. Microbiol. 159, 530–536.

Siponen M. I., Legrand P., Widdrat M., Jones S. R., Zhang W.-J., Chang M. C. Y., Faivre D., Arnoux P., Pignol D. (2013) Structural insight into magnetochrome-mediated magnetite biomineralization. Nature 502, 681–684.

Spangler B., Morgan C. W., Fontaine S. D., Vander Wal M. N., Chang C. J., Wells J. A., Rensio A. R. (2016) A reactivity-based probe of the intracellular labile ferrous iron pool. Nat. Chem. Biol. 12, 680–685.

Suzuki T., Okamura Y., Calugay R. J., Takeyama H., Matsunaga T. (2006) Global gene expression analysis of iron-inducible genes in *Magnetospirillum magneticum* AMB-1. J. Bacteriol. 188, 2275–2279.

Tang Y. Q., Dong Y. X., Wang X. F., Sriraghavan K., Wood J. K., Vennerstrom J. L. (2005) Dispiro-1,2,4-trioxane analogues of a prototype dispiro-1,2,4-trioxolane: mechanistic comparators for artemisinin in the context of reaction pathways with iron(II). J. Org. Chem. 70, 5103–5110.

Towe K. M., Bradley W. F. (1967) Mineralogical constitution of colloidal ‘hydrous ferric oxides”. J. Colloid. Interf. Sci. 24, 384–392.

Uebe R., Ahrens F., Stang J., Jäger K., Böttger L. H., Schmidt C., Matzanke B. F., Schüler D. (2019) Bacterioferritin of *Magnetospirillum gryphiswaldense* is a heterotetraeicosameric complex composed of functionally distinct subunits but is not involved in magnetite biomineralization. mBio 10, #e02795–18 doi: 10.1128/mBio.02795-18.

Uebe R., Schüler D. (2016) Magnetosome biogenesis in magnetotactic bacteria. Nature Rev. Microbiol. 14, 621–637.

Uebe R., Voigt B., Schweder T., Albrecht D., Katzmann E., Lang C., Böttger L., Matzanke B., Schüler D. Deletion of a *fur*-like gene affects iron homeostasis and magnetosome formation in *Magnetospirillum gryphiswaldense*. J. Bacteriol. 192, 4192–4204.

Wang Q., Wang X., Zhang W., Li X., Zhou Y., Li D., Wang Y., Tian J., Jiang W., Zhang Z., Peng Y., Wang L., Li Y., Li J. (2017) Physiological characteristics of *Magnetospirillum gryphiswaldense* MSR-1 that control cell growth under high-iron and low-oxygen conditions. Sci. Rep. 7: 2800. Doi: 10.1038/s41598-017-03012-4.

Wang X., Zhu M., Koopal L. K., Li W., Xu W., Liu F., Zhang J., Liu Q., Feng X., Sparks D. L. (2016) Effects of crystallite size on the structure and magnetism of ferrihydrite. Environ. Sci.: Nano. 3, 190–202.

Werckmann J., Cypriano J., Lefèvre C. T., Dembelé K., Ersen O., Bazylinski D. A., Lins U., Farina M. (2017) Localized iron accumulation precedes nucleation and growth of magnetite crystals in magnetotactic bacteria. Sci. Rep. 7, #8290 doi:10.1038/s41598-017-08994-9.

Zaitsev V. S., Filimonov D. S., Presnyakov I. A., Gambino R. J., Chu B. (1999) Physical and chemical properties of magnetite and magnetite-polymer nanoparticles and their colloidal dispersions. J. Colloid Interface Sci. 212, 49–57.

Zuoquan Z., Sun Q., Zhong J., Qihua Y., Li H., Cheng D., Liang B., Shuai X. (2014) Magnetic resonance imaging-visible and pH-sensitive polymeric micelles for tumor targeted drug delivery. J. Biomed. Nanotechnol. 10, 216–226.

